# DBF4, not DRF1, is the crucial regulator of CDC7 kinase at replication forks

**DOI:** 10.1101/2023.12.13.568031

**Authors:** Anja Göder, Chrystelle Maric, Michael D. Rainey, Chiara Cazzaniga, Daniel Shamavu, Jean-Charles Cadoret, Corrado Santocanale

## Abstract

In eukaryotes, CDC7 kinase is crucial for DNA replication initiation and has been involved in fork processing and replication stress response. Human CDC7 requires the binding of either one of two regulatory subunits, DBF4 and DRF1, for its activity. However, it is unclear whether the two regulatory subunits target CDC7 to a specific set of substrates, thus having different biological functions, or if they act redundantly.

Using genome editing technology, we generated an isogenic set of cell lines deficient in either one of the two CDC7-activating subunits: these cells are viable but present signs of genomic instability, indicating that both DBF4 and DRF1 can independently support CDC7 for bulk DNA replication. Nonetheless, DBF4-deficient cells show altered replication efficiency, including partial deficiency in MCM helicase phosphorylation and alterations in the replication timing of discrete genomic regions. Notably, we find that CDC7 function at replication forks is entirely dependent on DBF4 and not DRF1. Thus, DBF4 is the primary regulator of CDC7 activity, likely mediating most of its functions in unperturbed DNA replication and during replication fork processing upon replication interference.

## INTRODUCTION

In eukaryotic cells, DNA replication follows a highly regulated programme to ensure the faithful and complete duplication of the genome. This regulation is achieved through the spatio-temporal separation of origin firing and is linked to cellular mechanisms, including epigenome maintenance, gene expression and genome stability^1,2^.

An essential protein involved in the activation of replication origins is the Ser/Thr kinase CDC7, which phosphorylates multiple subunits of the MCM helicase complex at replication origins. This phosphorylation allows the recruitment of additional co-factors such as CDC45 and GINS and the activation of the helicase complex, allowing the unwinding of the double-stranded DNA for DNA replication^3–5^. CDC7-dependent phosphorylation of the MCM complex is counteracted by RIF1-PP1 phosphatase. Thus, the efficiency of replication initiation is determined by the interplay between kinases and phosphatases, contributing to the well-defined spatial-temporal coordination of origin firing^6,7^.

In addition to its role in origin activation, CDC7 kinase has important functions in the replication stress and DNA damage response. Upon replication stress, CDC7 facilitates the activation of the checkpoint kinase CHK1 through ATR by phosphorylating the mediator protein CLASPIN^8,9^. The ATR-CHK1 pathway is then responsible for determining cellular responses to replication stress, including transcriptional reprogramming and withdrawal from the cell cycle. Critically, the ATR-CHK1 pathway limits DNA damage and genome instability by stabilising stalled replication forks and preventing further origin firing^10,11^. Recently, we demonstrated that hCDC7 kinase participates in the processing of stalled forks by regulating the activity of the MRE11 nuclease, thus allowing more efficient fork restart and modulating fork speed upon mild replication stress. Partial CDC7 inhibition reduces the generation of ssDNA and fork collapse upon sustained fork arrest, which is mostly dependent on nuclease activity. Moreover, CDC7 physically associates with active and stalled replisomes, where it likely phosphorylates key substrates, possibly including MRE11 itself, as well as other proteins involved in homologous recombination DNA repair^12–14^.

In most eukaryotic organisms, CDC7 kinase activity is fully dependent upon its interaction with an activating subunit known as DBF4, which was first identified in budding yeast^15^, and the formation of a CDC7-DBF4 complex is thought to be the major mechanism controlling CDC7 activity. DBF4 is an evolutionarily conserved protein; however, sequence homology is restricted to three motifs, named N, M and C motifs, for their relative position in the protein. The M and C motifs interact with the N and C lobes of CDC7, respectively, and stabilise the kinase in an active conformation, while the N motif contains a BRCT-like domain and mediates protein-protein interactions^16–20^.

In human cells, two DBF4-like proteins have been functionally described: DBF4 and DBF4B. DBF4B is also called DRF1 (DBF4 Related Factor 1) or ASKL1^21,22^, but for clarity, we will refer to it as DRF1 throughout this work. Both proteins form stable complexes with CDC7; CDC7/DBF4 and CDC7/DRF1 complexes can be independently immunoprecipitated from cell extracts and have kinase activity in *in vitro* biochemical assays^23^. In human proliferating tissue and cancer cells, both DBF4 and DRF1 are expressed almost simultaneously, but in other species, their pattern of expression can greatly vary. In *Xenopus laevis*, DRF1 is only expressed during early embryonic development, where it is essential for DNA replication, however at later stages DRF1 expression is reduced and eventually replaced by DBF4^24,25^. On the other hand, *Saccharomyces cerevisiae* and *Drosophila melanogaster* only have one regulatory subunit, even though Drosophila expresses multiple variants of its DBF4 ortholog Chiffon^26,27^.

The concurrent expression of both subunits in human cells and their limited sequence homology raises the question of whether they each target CDC7 to a distinct set of substrates. Currently, very few targets of either DBF4 or DRF1 have been identified. DBF4 has been shown to specifically interact with the DNA topoisomerase TOP2A, whose activity at centromeres during S-phase is regulated by CDC7 phosphorylation^28^. Additionally, evidence has been provided of the CDC7-DBF4 complex being both a target and an important mediator of ATM/ATR-checkpoint signalling during replication stress and a promoter of trans-lesion DNA synthesis^29–31^. Our knowledge of DRF1, on the other hand, is comparably very limited, with most data coming from experiments in Xenopus egg extracts where DRF1 is the primary interaction partner of CDC7 and is essential for DNA synthesis^24,25^.

Even though the existence of two regulatory subunits for CDC7 has been known for over 20 years^21^, our progress in understanding DRF1 and DBF4’s contribution to CDC7 kinase activity in human cellular models has been limited due to technical challenges. Indeed, both endogenous proteins are extremely difficult to detect with antibody-based technologies because of their low levels of expression as well as the presence of multiple post-translational modifications, which possibly mask the epitopes recognised by most of the immunological reagents generated to date. While an early study indicated that siRNA depletion of DRF1 could delay S-phase and retard the growth of HeLa cells^32^, we consistently observed limited DRF1 reduction on mRNA level, even with multiple siRNAs.

In this work, we utilised genome editing to generate isogenic DBF4 and DRF1-deficient cells in MCF10A breast epithelial cells. Neither targeting DBF4 nor DRF1 compromises cell viability, indicating a partial redundancy between the two subunits to support the essential functions of CDC7 in DNA proliferation. However, DBF4 is the major contributor to CDC7 functions in DNA replication, and its deficiency causes alterations of the replication timing in discrete genomic regions similar to a partial inhibition of the CDC7 kinase with small molecules. Importantly, we identify DBF4 as the primary mediator of CDC7 activity during replication stress, where it contributes to fork processing and checkpoint signalling directly at stalled replication forks.

## RESULTS

### CDC7 activity is primarily mediated by DBF4 and only to a lesser extent by DRF1

To dissect the roles of CDC7’s regulatory subunits, we generated DBF4- and DRF1-deficient cell lines by transient transfection of MCF10A EditR cells^33^, stably expressing Cas9, with plasmids expressing short guide RNAs (sgRNAs) either targeting exon 3 of *DBF4* or exon 9 of *DRF1*. Both regions were explicitly chosen for their position upstream of the CDC7-activating M and C motifs (Figure 1A-B). From the transfected pools, two independent clones that had undergone gene editing were isolated. For DBF4, MCF10A DBF4 clone 11 and 30 (DBF4-11 and DBF4-30) displayed homozygous deletions of 13 and 5 nucleotides, respectively. For DRF1, two monoclonal cell lines were generated, MCF10A DRF1 clone 5 (DRF1-5) with a homozygous 4 nucleotides deletion and MCF10A DRF1 clone 7 (DRF1-7), with a single nucleotide insertion into exon 9 of *DRF1* (Figure S1). In all cases, gene editing resulted in a premature stop codon, generating truncated proteins lacking the critical domains required for the binding and activation of CDC7, which could be considered a bona fide loss of function with respect to CDC7 activation. As a note of caution, it is not possible to fully exclude that through mRNA translation from an internal start site or exon skipping events, low levels of proteins still capable of providing some functionality may be produced; thus, throughout this work, we define these cell lines as DFB4- and DRF1-deficient and not knockout cells. If not otherwise indicated, experiments were preferentially performed with MCF10A DBF4-11 and DRF1-7 cells.

**Figure 1:**
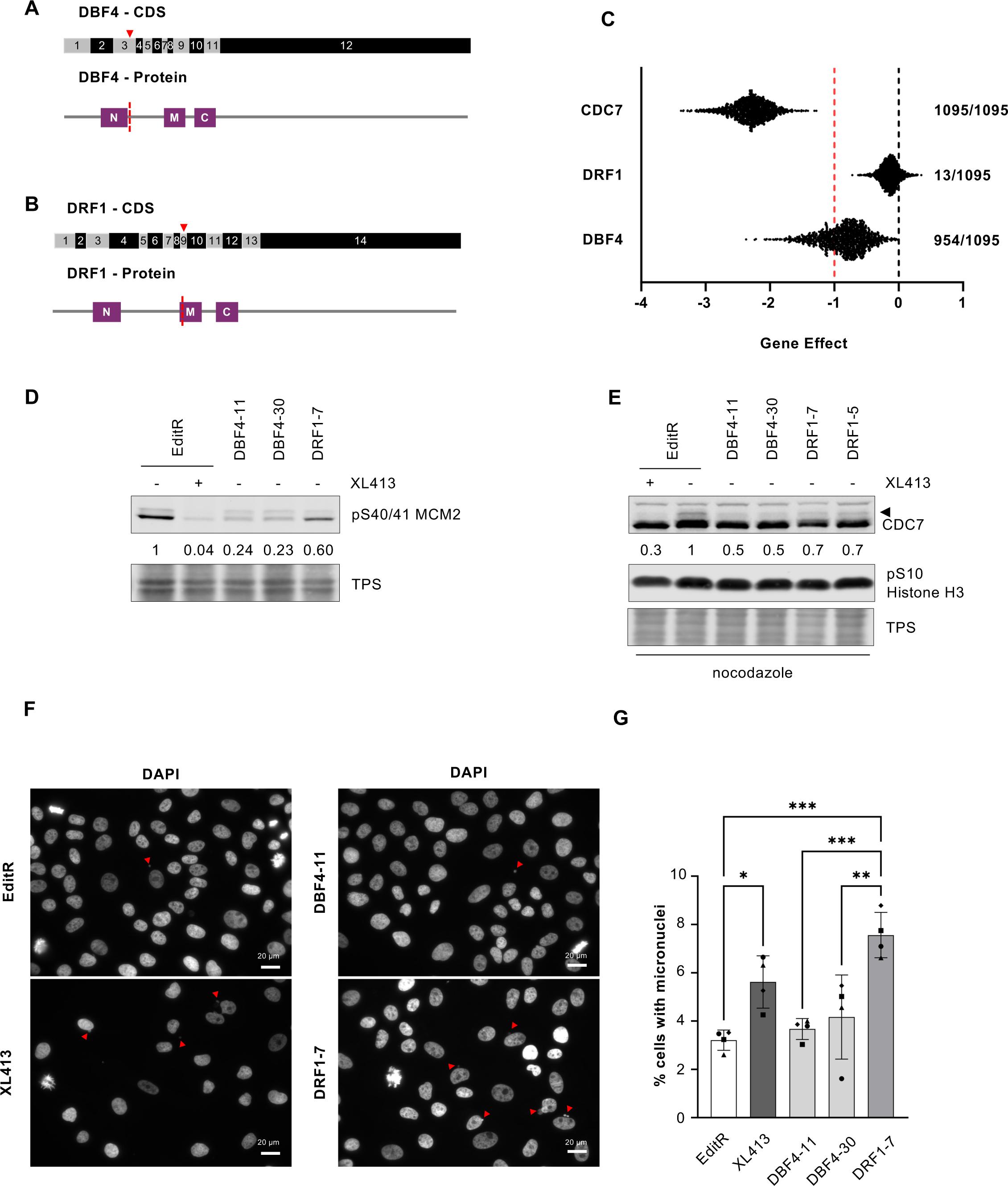
CDC7 activity is primarily mediated by DBF4 and only to a minimal extend by DRF1. **A-B.** Schematic representation of gene editing approach for the generation of DBF4-**(A)** and DRF1-deficient **(B)** cells. Red triangle marks the position of the Cas9 cut site in the coding sequence (CDS) for DBF4 or DRF1. Protein of both subunits displays relevant motifs (M, N, and C motif) aligned with the CDS and marks the position of the Cas9 cut site relative to the protein sequence. **C.** Comparison of CHRONOS dependency scores (Gene effect) obtained in CRISPR-Cas9 knockout screens for CDC7, DBF4 and DRF1 available from DepMap portal. A score of 0 (black line) or higher would describe a non-essential gene, a score of −1 (red line) corresponds to the median of all pan essential genes. A lower score describes a higher chance of the gene of interest being essential. Numbers represent the number of cell lines, which have been classified as dependent on CDC7, DBF4 or DRF1 of 1095 tested cell lines. **D.** MCF10A EditR, MCF10A DBF4-11 and −30, and MCF10A DRF1-7 were treated with 10 µM XL413 or DMSO for 24 hrs. Whole cell extracts were prepared and analysed by immunoblotting with indicated antibodies. Total protein staining (TPS) was used as loading control. Numbers indicate relative changes in MCM2 phosphorylation compared to the DMSO-treated control cell line and normalised to TPS in the displayed blot. Data are representative of at least three independent experiments. **E.** MCF10A EditR, MCF10A DBF4-11 and −30, and MCF10A DRF1-5 and −7 were treated with 10 µM XL413 or DMSO for 24 hrs. 16 hrs before harvesting cells were additionally treated with 0.2 µg/ml nocodazole. Whole cell extracts were prepared and analysed by immunoblotting with indicated antibodies. Total protein staining (TPS) was used as loading control. Triangle marks the mobility shift of CDC7 and numbers indicate relative changes in CDC7 mobility shift compared to the DMSO-treated control cell line and normalised to TPS of the displayed blot. Data are representative of three independent experiments. **F.** MCF10A EditR, were either mock or treated with 10 µM XL413 for 24 hrs. MCF10A DBF4-11 and −30, and MCF10A DRF1-7 were only mock-treated. Cells were fixed, stained with DAPI to visualise DNA, and analysed by fluorescence microscopy. Representative images of four independent experiments are shown (Scale bar, 20 µm). Red triangles indicate micronuclei. Brightness of images was adjusted for all samples to aid visualisation. **G.** Graph shows percentage of cells with micronuclei for four independent experiments, mean ± SD. At least 275 cells were analysed per condition for each experiment. Statistical analysis was performed using one-way ANOVA with Tukey’s multiple comparison test (*p<0.05, **p<0.01, ***p<0.001).

The generation of viable DBF4- and DRF1-deficient cell lines indicates that either DBF4 or DRF1 can support CDC7’s essential function in cell proliferation. Our attempts to knockout both *DBF4* and *DRF1* were unsuccessful, as we were unable to recover viable clones, similar to our previous attempts at generating a CDC7 knockout ^33^, further providing evidence for partial redundancy between DBF4 and DRF1. This result indicates that, unlike CDC7, neither DBF4 nor DRF1 is essential in MCF10A cells and led us to inspect the datasets from multiple CRISPR/Cas9 screens, which are publicly available on DepMap Portal, assessing the dependency of multiple cell lines on a given gene for proliferation^34,35^. All the 1095 cell lines tested showed a very strong dependency on CDC7, therefore defining CDC7 as a common essential gene. Differently, most tested cell lines displayed some level of dependency on DBF4, but this was less marked than on CDC7, while only 13 of 1095 cell lines showed a dependency on DRF1 (Figure 1C). Altogether, these findings indicate that the two CDC7 activating subunits differently contribute to cell proliferation in human cells.

To assess DBF4 and DRF1 contributions to the overall CDC7 activity in a cellular context, we analysed the levels phosphorylation of the helicase subunit MCM2, a well-established CDC7 substrate^36^, in MCF10A EditR, DBF4- and DRF1-deficient cells, using an antibody that recognises MCM2 when phosphorylated at both S40 and S41. As a control, MCF10A EditR were treated with the CDC7 inhibitor (CDC7i) XL413. XL413 treatment nearly completely abolished MCM2 phosphorylation, while the loss of DBF4 drastically reduced it. In comparison, MCF10A DRF1-7 only showed a mild reduction of pS40/S41 MCM2 (Figure 1D). As CDC7-dependent phosphorylation of MCM2 at S40 is cell cycle regulated and dependent on a priming kinase phosphorylating S41^36^, we also assessed CDC7 phosphorylation in cells arrested in mitosis. Under these conditions, CDC7 has been shown to be phosphorylated at multiple residues, some of which are potentially due to autophosphorylation, resulting in an electrophoretic mobility shift during SDS-PAGE^37,38^. MCF10A EditR, MCF10A DBF4- and DRF1-deficient cells were arrested in mitosis using nocodazole, an inhibitor of microtubule polymerisation, and then protein extracts were prepared and analysed by immunoblotting. We observed that in parental cells, a fraction of CDC7 molecules displayed altered electrophoretic mobility upon nocodazole treatment, which was strongly attenuated by treatment with XL413, thus demonstrating that the change in mobility depends on CDC7 activity. Both the loss of DBF4 and DRF1 reduced the mobility shift of CDC7 by approximately 50% and 30%, respectively (Figure 1E).

CDC7 inhibition is associated with irregular progression through mitosis, often resulting in the formation of micronucleated cells^39,40^. Interestingly, while we confirmed that the treatment with XL413 increased the percentage of cells with micronuclei, we did not observe a significant change in the percentage of micronucleated cells in DBF4-deficient cells compared to parental cells. However, micronucleated cells accumulated in DRF1-deficient cells (Figure 1F-G).

Taken together, these experiments indicate that both DBF4 and DRF1 can support the fundamental functions of CDC7 kinase in cell proliferation and that in MCF10A cells, DBF4 is the main contributor to overall CDC7 activity, thus suggesting a potential and possibly more restricted role of DRF1.

### DBF4, not DRF1, is the major contributor to CDC7 activity in DNA replication

To assess if DBF4 and DRF1 deficiency directly affect the dynamics of DNA replication, we performed a flow cytometry-based analysis of DNA synthesis by pulse labelling parental, DBF4- and DRF1-deficient cells for 30 minutes with the thymidine analogue EdU. We noticed that EdU incorporation in nascent DNA was only partially reduced in DBF4-deficient cells, an effect that was more evident in cells in mid to late S-phase. Interestingly, this effect was not as pronounced as the direct inhibition of the kinase with XL413 and, more importantly, was not observed in DRF1-deficient cells (Figure 2A-B). Consistent with the results obtained with engineered cell lines, siRNA targeting DBF4 reduced the rate of DNA synthesis in MCF10A EditR cells, while the transfection of DRF1-targeting siRNAs did not affect DNA synthesis (Figure S2A-D). By targeting both subunits in parental cells with siRNA, we did not further reduce the rate of EdU incorporation compared to siDBF4 alone (Figure S2C-D). This result was further confirmed by the lack of reduction in the rate of DNA synthesis when transfecting siRNA targeting DRF1 into DBF4-deficient cells, whereas targeting DBF4 in MCF10A DRF1-deficient cells reduced but did not abolish EdU incorporation (Figure S2D-E). Downregulation of the DBF4 and DRF1 mRNA was monitored by qPCR, and while DBF4 siRNAs efficiently downregulated DBF4 expression, the efficiency of DRF1 siRNA was limited to approximately 60% with residual expression likely masking relevant phenotypes (Figure S2A-B).

**Figure 2:**
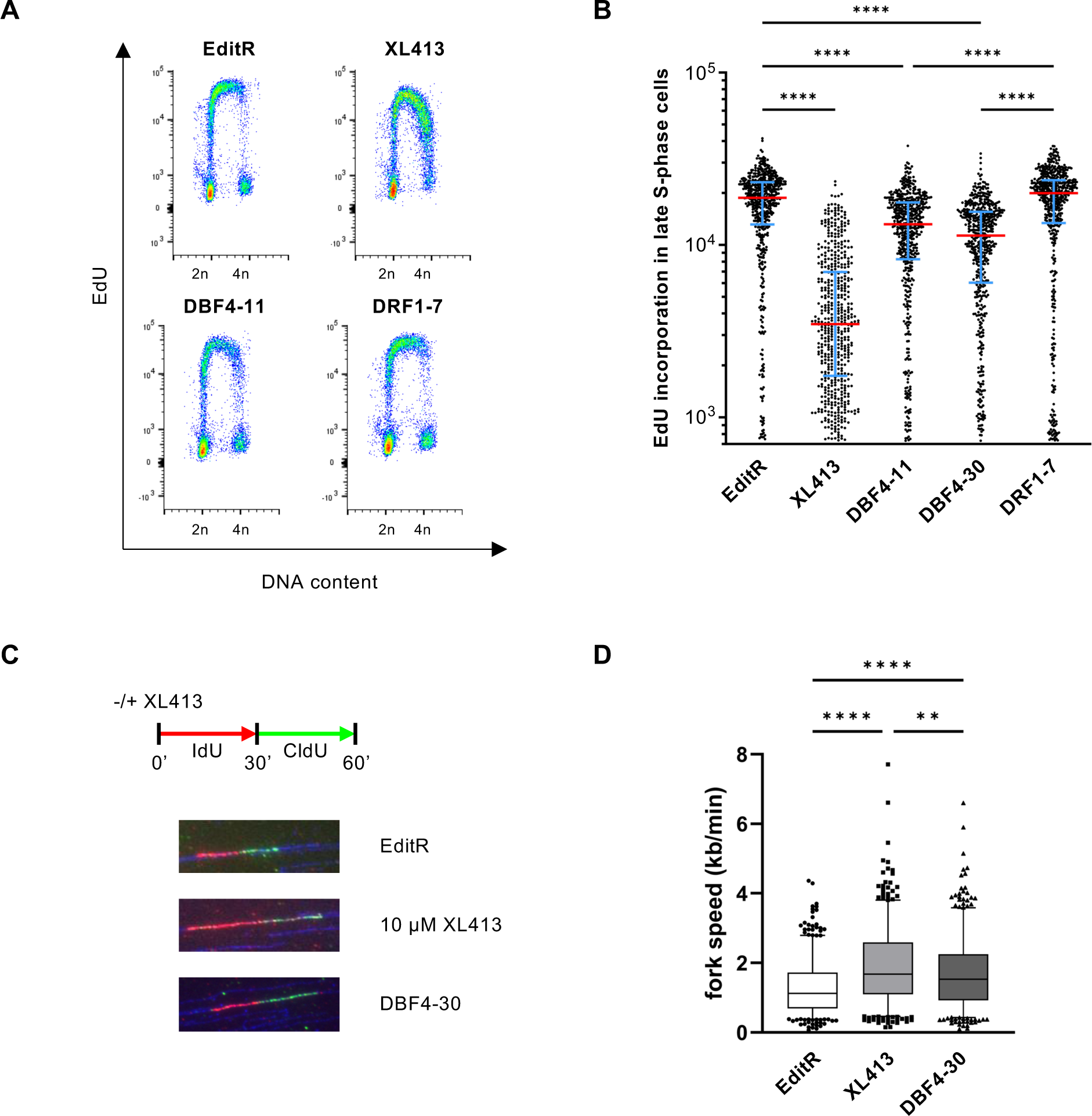
DBF4, not DRF1, is the major contributor to CDC7 activity in DNA replication. **A.** MCF10A EditR, MCF10A DBF4-11 and −30, and MCF10A DRF1-7 were treated with 10 µM XL413 or DMSO for 24 hrs. For flow cytometry analysis, cells were labelled with 10 µM EdU 30 min prior to harvest. Representative images from one of three independent experiments are shown. **B.** Fluorescence intensity, proportional to EdU incorporation, in late S-phase cells for representative experiment displayed in **A**. Red lines indicate the medians and blue lines show the interquartile range extending from the 25^th^ to the 75^th^ percentile of 552 cells. Statistical analysis was performed using one-way ANOVA with Tukey’s multiple comparison test (****p<0.0001). **C.** MCF10A EditR and MCF10A DBF4-30 were treated with 10 µM XL413 or DMSO for 24 hrs and then labelled with IdU (red) for 30 min. IdU was washed off and cells were labelled with CldU for 30 min in the presence of 10 µM XL413 or DMSO. Representative fibres for each treatment/cell line are shown. **D.** Analysis of replication fork speed in the experiment described in **C**. At least 100 tracks were analysed for each condition of three independent experiments. Box plots display the median and 5-95% range. Statistical analysis was performed using one-way ANOVA with Tukey’s multiple comparison test (**p<0.01, ****p<0.0001).

Since these experiments did not reveal a direct involvement of DRF1 in DNA synthesis and suggested that DBF4 could be the primary activator of CDC7 in DNA replication, we further investigated the DNA replication dynamics in parental and DBF4-deficient cells by DNA combing assay. We found that in DBF4-deficient cells, the average replication fork speed was increased from 1.2 kb/min to 1.7 kb/min (Figure 2C-D). This phenotype is most likely dependent on the decrease in overall CDC7 activity and can be recapitulated by partial inhibition of the kinase with XL413 (Figure 2C-D). Such observations are consistent with a partial loss of CDC7 activity limiting origin firing, which is compensated for by an increase in replication fork speed, as previously determined with other CDC7 small molecule inhibitors^41,42^.

### Loss of DBF4 changes global replication timing similar to CDC7 inhibition

To further characterise how CDC7 and DBF4 regulate DNA replication at a genome-wide level, we performed replication timing experiments in MCF10A EditR with or without CDC7i treatment and in the isogenic DBF4-deficient cells. In this series of experiments, cells were labelled with a short pulse of BrdU and divided into early and late S-phase fractions. The neo-synthesized DNA was hybridised on human whole genome microarrays, as previously described^43^. Replication timing differential analyses on MCF10A EditR cells and either CDC7i-treated or DBF4-deficient cells were performed using the START-R suite software^44^ and visualised for the whole genome. The comparison of the profiles of MCF10A EditR with either CDC7i-treated or DBF4-deficient cells showed variations in the replication timing of multiple regions across the whole genome (Figure 3A-B, Figure S3A-B). Altered replication timing regions are mostly distributed in clusters of several adjacent advanced or delayed regions (see illustration for chr1 in Figure 3A-B). In XL413-treated MCF10A, 20.6% of the genome showed an altered replication timing programme compared to the MCF10A EditR cells, whereas 13.7% of the genome is altered in DBF4-deficient cells. Globally, 322 regional changes in replication timing were identified after CDC7i treatment, with 125 regions showing advanced and 197 showing delayed timing. Interestingly, the loss of DBF4 led to 185 changes in replication timing, 58 of those showing advanced and 127 regions showing delayed timing. In line with DBF4 being required for most of CDC7’s activity, approximately 70% of changing regions in DBF4-deficient cells were similarly affected by CDC7 inhibition (Figure 3C). We also noted that around 2/3 of the perturbations of the replication timing at the genome scale resulted in delayed genomic regions either after CDC7 inhibition or DBF4 loss. Advanced regions in XL413-treated cells and DBF4-deficient cells mainly were regions that were initially early replicating regions; however, some late replicating regions and TTR (timing transition regions) also displayed earlier timing than in the control cell line (Figure 3A-B, 3D-E, and Figure S3A). Similarly, delayed regions for both samples could be primarily found in parts of the genome replicating in early S-phase or in TTR regions (Figure 3A-B, 3D-E and Figure 3SB). A more in-depth look at potential commonalities between the delayed or advanced regions revealed that delays in replication timing were often found in large genes (>400 kb) (Figure S3C), which are known to be poor in replication origins and prone to replication stress^28^. In line with this, delayed regions for both XL413-treated and DBF4-deficient cells were poor in putative G4 sequences and CpG islands, which are often enriched at replication origins^45^ and rarely contained constitutive origins (Figure S3D-F), suggesting that these regions are likely to be replicated passively. However, advanced regions in XL413-treated cells and DBF4-deficient cells were largely enriched in CpG islands and putative G4 sequences as well as constitutive origins (Figure S3D-F), even more so than regions traditionally known to replicate in early S-phase. This likely enables those regions to replicate despite the perturbation in origin firing by CDC7 inhibition. Taken together, DBF4 deficiency alters the replication timing in discrete genomic regions in a similar manner to the treatment with the CDC7 inhibitor XL413.

**Figure 3:**
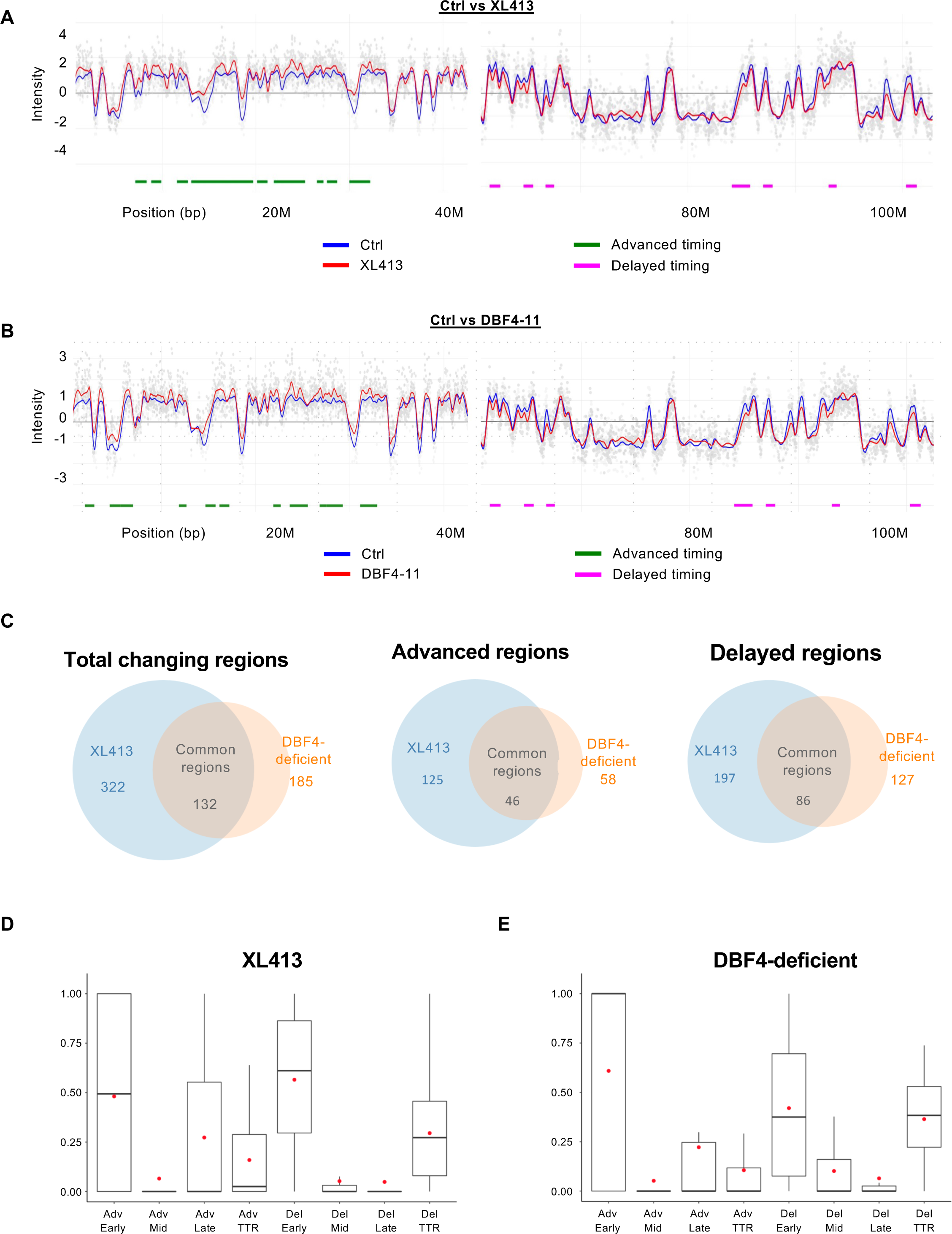
Loss of DBF4 and CDC7 inhibition induce similar changes in global replication timing. MCF10A EditR and MCF10A DBF4-11 were treated with 10 µM XL413 or DMSO for 24 hrs before replication timing analysis. **A.** Part of chromosome 1 replication timing profiles in untreated or XL413-treated MCF10A EditR cells cells. Blue line represents MCF10A EditR cells treated with DMSO, the red one cells treated with 10 µM XL413. Chromosome coordinates are indicated below the profile in megabases. Differences in replication timing are marked below the profile with advanced regions in green and delayed regions in magenta. Data is representative of two replicates of four independent experiments. **B.** Part of chromosome 1 replication timing profiles for MCF10A EditR compared to MCF10A DBF4-deficient cells (DBF4-11). Analysis was performed and graphs were generated as described in **A**. **C.** Summary of changes in replication timing in MCF10A EditR treated with 10 µM XL413 (blue) and MCF10A DBF4-deficient cells (orange) displayed as a Venn diagram for total changing regions (left), advanced regions (middle), and delayed regions (right). Numbers represent the numbers of changed regions for indicated samples. Four independent experiments were performed for XL413-treated MCF10A cells and for DBF4 deficient cells. **D.** Analysis of replication timing changes in MCF10A EditR treated with 10 µM XL413 for 24 hrs relative to the replication timing regions they originated from; either Early, Mid, and Late replicating regions or Timing Transition Regions (TTR). Advanced regions (Adv) and Delayed regions (Del) are displayed separately. The box plots show the dispersion of the data with a range from the 25^th^ to 75^th^ percentile, the sample median is represented by the line inside the box and the mean by a red dot. **E.** Analysis of replication timing changes in MCF10A DBF4-deficient cells treated with DMSO for 24 hrs as described in **D**.

### DBF4 mediates the majority of CDC7 functions in the replication stress response

We previously reported that CDC7, independent of its role in origin firing, promotes the processing and collapse of stalled replication forks upon prolonged treatment with hydroxyurea (HU) and that inhibition of CDC7 activity results in suppression of checkpoint signalling and DNA double-strand break (DSB) induction^12^. To understand whether a specific regulatory subunit mediates this role of CDC7, we treated MCF10A EditR, MCF10A DBF4- and DRF1-deficient cell lines with 4 mM HU for 16 hrs and examined the induction of the replication stress response via protein fractionation and Western blot analysis. As a control, MCF10A EditR cells were treated with 10 µM XL413 30 min prior to HU treatment (Figure 4A-B). The inhibition of CDC7 with XL413, as well as the loss of DBF4, led to a reduction of CHK1 phosphorylation at Ser345, a downstream marker of ATR activity, while in DRF1-deficient cells, CHK1 phosphorylation was not compromised (Figure 4A). Interestingly, even though CHK1 activation was affected by CDC7 inhibition and mutation of DBF4, the autophosphorylation of ATR itself was not obviously reduced (Figure 4B), suggesting that DBF4 with CDC7 mediates an intermediate step in signalling amplification, likely the well-characterised CDC7-dependent phosphorylation of CLASPIN which facilitates CHK1 activation by ATR^8,46,47^.

**Figure 4:**
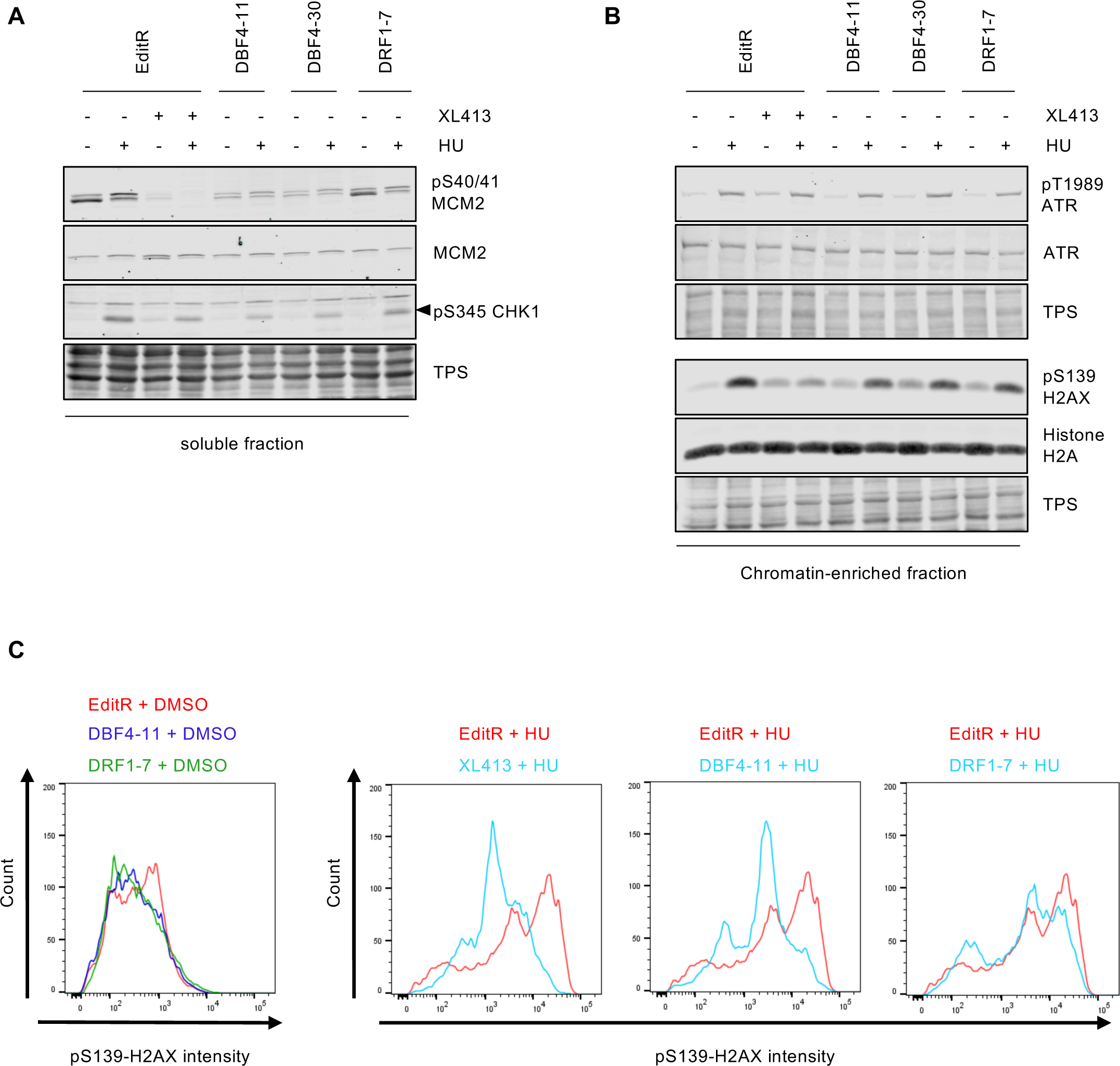
DBF4 mediates CDC7 activity in the replication stress response. **A-B.** MCF10A EditR, MCF10A DBF4-11 and −30, and MCF10A DRF1-7 were pre-treated with 10 µM XL413 or DMSO for 30 min before the addition of 4 mM HU for 16 hrs. Soluble (**A**) and chromatin-enriched (**B**) fractions were prepared and analysed by immunoblotting with indicated antibodies. TPS was used as loading control. Data are representative of three independent experiments. **C.** MCF10A EditR, MCF10A DBF4-11 and −30, and MCF10A DRF1-7 were pre-treated with 10 µM XL413 or DMSO for 30 min before the addition of 4 mM HU for 24 hrs. For flow cytometry analysis of pS139-H2AX intensity, cells were harvested fixed and stained with the indicated antibody. Representative images from one of three independent experiments are shown. MCF10A EditR treated with HU in the panels is the same sample in pairwise comparison with other samples for better visualisation.

Replication fork stalling induces the phosphorylation of the histone variant H2AX at Ser139 (γH2AX), which is further increased by replication fork collapse and DSB formation^48^. As previously described by western blotting, we observed that inhibition of CDC7 suppresses the phosphorylation of H2AX upon HU treatment^12^. Intriguingly, H2AX phosphorylation also appeared to be partially reduced in both DBF4- and DRF1-deficient cells (Figure 4B). To accurately quantify the contribution of DBF4 and DRF1, we returned to a quantitative flow cytometry-based assay to analyse γH2AX phosphorylation levels in single cells. Following a 24-hour treatment with HU, MCF10A EditR cells showed a clear induction of γH2AX, which was drastically reduced by pre-treatment XL413 or in DBF4-deficient cells, while the reduction in H2AX phosphorylation in DRF1-deficient cells was much more limited (Figure 4C). Taken together, the results indicate that DBF4 is the primary mediator of CDC7 activity in promoting H2AX phosphorylation upon prolonged fork stalling, whereas DRF1 has, at most, a minor role in this process.

### CDC7 activity at stalled replication forks is mediated by DBF4

We previously described that CDC7 promotes MRE11-dependent processing of stalled replication forks and we identified a small, discrete pool of MRE11, that displays a CDC7-dependent, phosphatase-sensitive electrophoretic mobility shift suggesting a CDC7-dependent phosphorylation event^12^.

To determine if MRE11 phosphorylation is dependent on either CDC7-DBF4 or CDC7-DRF1 complexes, we prepared protein extract from MCF10A EditR, either mock or treated with XL413, MCF10A DBF4-11 and DBF4-30 as well as MCF10A DRF1-7 cells treated with 4 mM HU for 24 hrs. MRE11 showed a clear mobility shift in HU-treated MCF10A EditR cells, which, as expected, was drastically reduced in samples treated with CDC7i. In line with previous results, the DBF4 deficiency substantially decreased the mobility shift of MRE11, while the mutation of DRF1 only showed a limited reduction (Figure 5A and S4). To assess the impact of DBF4 and DRF1 deficiency on CDC7 functions directly at replication forks, we extracted nascent DNA and analysed associated proteins by DNA-mediated chromatin pull-down (Dm-ChP)^49^. In a first set of experiments, we found that CDC7 was recruited at replication forks in an XL413-independent manner, and it was present in both DBF4- and DRF1-deficient cells during unperturbed DNA replication (Figure 5B). We then treated MCF10A EditR and MCF10A DBF4-11 cells with 4 mM HU over 24 hours and observed that the mobility shift of MRE11 at the stalled replication forks was lost in both XL413-treated and DBF4-deficient cells correlating with the suppression of H2AX and RPA2 phosphorylation (Figure 5C). In a parallel experiment using DRF1-deficient cells instead, MRE11 phospho-shift, H2AX and RPA2 phosphorylation were either not affected or only very partially compromised (Figure 5D), thus identifying DBF4 as the regulatory subunit involved in CDC7’s function at forks.

**Figure 5:**
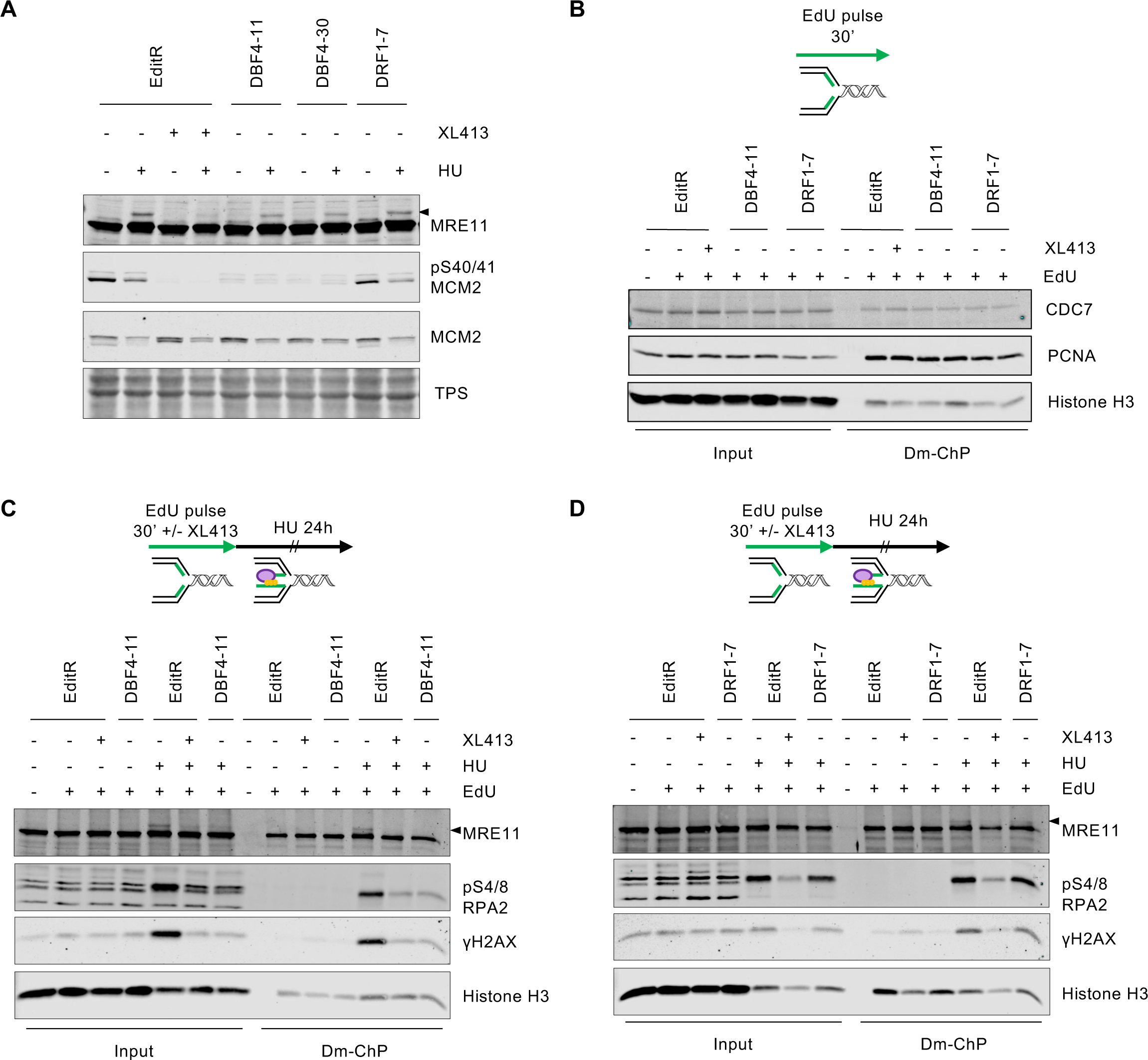
CDC7’s activity at replication forks is solely mediated by DBF4. **A.** MCF10A EditR, MCF10A DBF4-11 and −30, and MCF10A DRF1-7were pre-treated with 10 µM XL413 or DMSO for 30 min before the addition of 4 mM HU for 24 hrs. Whole cell extracts were prepared and analysed by immunoblotting with indicated antibodies. Total protein staining (TPS) was used as loading control. Triangle marks MRE11 electrophoretic mobility shift. Data are representative of three independent experiments. **B.** MCF10A EditR, MCF10A DBF4-11, and MCF10A DRF1-7 were treated with 10 µM XL413 or DMSO for 24 hrs. Before harvesting cells were labelled with EdU for 30 min, then cells were fixed and proteins binding to EdU-labelled DNA captured by the DNA-mediated chromatin pull-down technique (Dm-ChP). Graphical representation of the experiment is shown above the Western blot analysis. Both input and captured material (Dm-ChP) were analysed by Western blot with indicated antibodies. Histone H3 was used as a loading control. Unlabelled cells (-EdU) were used as a negative control for the Dm-ChP samples. Data are representative of two independent experiments. **C.** MCF10A EditR and MCF10A DBF4-11were pre-treated with 10 µM XL413 or DMSO and 10 µM EdU for 30 min followed by a treatment with 4 mM HU for 24 hrs. Samples were prepared as described in **B**. Data are representative of three independent experiments. **D.** MCF10A EditR and MCF10A DRF1-7 were pre-treated with 10 µM XL413 or DMSO and 10 µM EdU for 30 min followed by a treatment with 4 mM HU for 24 hrs. Samples were prepared as described in **B**. Data are representative of three independent experiments.

## DISCUSSION

The functions of CDC7 have been primarily investigated without distinction between CDC7-DBF4 and CDC7-DRF1 complexes and whether they target CDC7 to a distinct set of substrates or if both regulatory subunits can support the various functions of the kinase. *In vitro* experiments have shown that both subunits are capable of activating CDC7 kinase and mediating the phosphorylation of the MCM complex^21^, which is likely due to the presence of the CDC7 binding motifs M and C in both subunits. Similarly, the switch from DRF1 to DBF4 in *Xenopus laevis* over the course of its development^50,51^ suggests that there is at least some redundancy between the two subunits in supporting the essential activity of CDC7 in DNA replication in vertebrates. In line with this, our data show that cells with a deficiency in either subunit are viable and able to proliferate, while we were unable to produce a double knockout of DBF4 and DRF1.

However, despite the basic redundancy between DBF4 and DRF1, our results indicate in MCF10A cells a preference for DBF4 when it comes to mediating CDC7 activity in DNA replication and the replication stress response. Such a preference for DBF4 may be either quantitative, i.e. higher DBF4 than DRF1 protein levels in cells, or due to different structural properties affecting the strength of key protein-protein interactions, thus driving CDC7 to its targets with different affinities.

Firstly, we demonstrate that the loss of DBF4 has a detectable impact on the rate of DNA synthesis, which goes hand in hand with an increase in replication fork speed. This can likely be attributed to a reduction in origin firing since it forces DNA replication forks to move faster to compensate for the reduced number of active forks^52^. This is further underlined by the changes observed in replication timing, which favour a delay in timing in regions that are sparse in origins, low on CpG islands and putative G4, and therefore rely on passive replication. In contrast, regions with a high density of origins have higher chances of being activated even if CDC7 activity is partially compromised. A recent study showed that severe inhibition CDC7 leads to S-phase blockade with partially replicated DNA, mostly in late replicating domains^13^. In our study, where CDC7 inhibition is not as stringent and genome duplication can be completed, we observed an advancement in the replication timing in specific regions within late replicating domains. Interestingly, we also detected a delay in replication timing for some early replicating regions, suggesting that in some genomic regions we potentially have an early-to-late and late-to-early shift in the replication timing programme. We found that the common regions of advanced and delayed timing for CDC7 inhibition and DBF4 loss showed mostly boundary shifts, events also frequently observed by Hiratani and collaborators during mouse embryonic cells differentiation^53^.

Major changes in replication time have been described in Rif1-deficient cells, a protein that contributes to nuclear architecture, DNA repair and that, in replication, counteracts the phosphorylation of the MCM complex by CDC7^6,54^. Such changes are strongly cell type dependent and are reinforced upon several rounds of replication, correlating with redistribution of active and inactive chromatin marks and alteration in chromatin architecture^55^. While it is difficult to directly compare replication timing (RT) profiles generated in different experimental models, we observe some analogies between our results and the ones obtained in mouse embryonic stem cells after the loss of RIF1-PP1 interaction^56^. In DBF4-deficient and CDC7-inhibited cells and similarly, in RIF-PP1-deficient cells, the RT profiles show a better degree of distinction between Early and Late replicating domains than in RIF1-KO, where the timing program is distributed solely toward mid S-phase regions^55^. This could raise the possibility that protracted CDC7 inhibition may lead to epigenetic perturbation, a hypothesis that should be experimentally tested in future studies.

Secondly, the functions of CDC7 in replication checkpoint signalling and replication fork processing are mediated by DBF4. While the activation of ATR upon drug-induced replication stress is still effective in DBF4-deficient and CDC7-inhibited cells, the downstream signalling is impaired. The lack of CHK1 phosphorylation is likely attributable to CDC7’s phosphorylation of CLASPIN which has been shown to be required for effective activation of CHK1^8^ and, considering our results, is most likely mediated by DBF4. On the other hand, the suppression of H2AX and RPA2 phosphorylation at stalled replication forks cannot be solely attributed to the lack of CHK1 activation but is most likely dependent on reduced MRE11-dependent processing and degradation of stalled replication forks; this is consistent with defective MRE11 phosphorylation in DBF4-deficient cells. Intriguingly, while CDC7’s function at forks requires DBF4, its recruitment is unaffected by DBF4 or DRF1 deficiency. At this stage, we can speculate that CDC7 may be recruited to the replisome by protein-protein interactions mediated by either DBF4 and DRF1, with only DBF4 being able to target the kinase to key substrates, or that its recruitment is solely dependent on domains located on CDC7.

Still, enigmatically, DRF1, while able to support the essential CDC7 activity in cell division, shows very limited involvement in mediating known specific functions of CDC7 compared to its counterpart DBF4. An increased number of micronucleated cells is the main phenotype we observed in DRF1-deficient cells, which could be consistent with a specific but not yet identified function in chromosome segregation, in the fine-tuning of DNA replication or the DNA repair process.

Expanding our knowledge of CDC7’s biological role and how these are driven by its regulatory subunits will be critical for the rational development of CDC7 inhibitors and CDC7-targeted therapies. The overexpression of CDC7 in multiple cancer types is often accompanied by an increased expression of DBF4^57^. This is in line with the importance of DBF4 for both efficient DNA replication and the replication stress response we demonstrated here, two processes that cancer cells rely on heavily.

Attempts at targeting CDC7 with small molecule ATP competitor inhibitors, while very effective in preclinical models, have so far shown limited success in clinical trials. With new technologies emerging capable of specifically affecting non-enzymatic proteins, such as “molecular glues” and genome editing approaches, it is tempting to suggest that the direct targeting of DBF4 may result in increased therapeutic window by avoiding the blockade of DNA replication in normal cells, thus reducing the potential for toxicity, and in the identification of cancer types specifically dependent on DBF4 expression and function.

## ACKNOWLEDGEMENTS

We thank Raimundo Freire for his help and commitment in the attempts to generate anti-DRF1 antibodies. We thank Enda O’Connell at the screening and genomics core and Shirley Hanley at Flow Cytometry Core Facility at University of Galway, which is supported by University of Galway, Science Foundation Ireland, the Irish Government’s Programme for Research in Third Level Institutions, Cycle 5 (PRTLI5) and the European Regional Development Fund. We also thank Nicolas Valentin for performing the flow cytometry cell sorting at the ImagoSeine core facility of the Institut Jacques-Monod.

This work was supported by Science Foundation Ireland (SFI) grant 16/IA/4476 and by La Ligue Nationale Contre le Cancer (RS16/75-108 and RS17/75-135), the GEFLUC, the Institut National du Cancer INCa-10493, the IdEx Université de Paris ANR-18-IDEX-0001 and by the generous legacy of Ms. Suzanne Larzat to Jean-Charles Cadoret’s group. Daniel Shamavu is supported by a Government of Ireland Postgraduate Scholarship the Irish Research Council under grant number GOIPG/2022/896.

## Author’s contribution

Conceptualization, AG, CM, MDR, JCC and CS; Investigation AG, CM, MDR, CC and DS; Writing – Original Draft AG, CM and CS; Writing – Review & Editing AG, CM and CS; Supervision JCC and CS; Funding Acquisition JCC and CS.

## METHOD DETAILS

### Cell culture

Cell culture was performed in a Class II Bio-safety cabinet, and all cell lines were maintained at 37°C in a humidified atmosphere containing 5% CO2. Cell counts and viability were determined using trypan blue exclusion and a countess (Invitrogen) or LUNA II cell counter (Logos biosystems). MCF10A cell line (human, female origin) was purchased from ATCC and authenticated via whole genome sequencing. MCF10A EditR cells (stably expressing Cas9) were previously described^33^. MCF10A DBF4-deficient clones 11 and 30, as well as DRF1-deficient clones 5 and 7, were derived via CRISPR/Cas9 genome editing from MCF10A EditR cells and monoclonal expansion. The mutation of DBF4 or DRF1 was verified via targeted PCR and Sanger sequencing. MCF10A cells and derivatives were cultured using DMEM supplemented with 5% (v/v) horse serum, 25 ng/ml cholera toxin, 10 μg/ml insulin, 20 ng/ml epidermal growth factor (Peprotech), 500 ng/ml hydrocortisone, 50 U/ml penicillin and 50 μg/ml streptomycin. Lenti-X™ 293T cell line (human, female origin) were purchased from TaKaRa and used without further authentication. Cell culture was performed using DMEM +GlutMAXTM-I (ThermoFisher) supplemented with 10% (v/v) fetal bovine serum, 50 U/ml penicillin and 50 μg/ml streptomycin. Unless otherwise specified, all culture media and reagents were obtained from Merck.

### Drugs treatments

If not otherwise indicated, XL413 (synthesised in-house) was used at a concentration of 10 μM. Hydroxyurea (HU) was used at 4 mM and nocodazole at 0.2 µg/ml (all acquired from Merck).

### DNA transfections

Cells were transfected with plasmid DNA (0.75 µg) using a 1:3 (w/v) ratio of DNA to polyethyleneimine “MAX” MW 40,000 (1 mg/ml, Polysciences). These were first individually diluted in 100 µl of 150 mM NaCl, mixed and incubated at room temperature for 20 min before being added to the cells.

### Generation of DBF4- and DRF1-deficient monoclonal cell lines

Oligonucleotides coding for sgRNAs targeting *DBF4* (5’-TGGGTCGAATTTCTCCTGTA-3’) or *DRF1* (5’-CGTTCCTCAAAATCGAAGAT-3’) were cloned into Bbs1 restriction site of the pX330-Hygromycin plasmid as previously described^33^. MCF10A EditR cells were transiently transfected with pX330-Hygromycin vectors carrying DBF4-sgRNA and DRF1-sgRNA. Cells were cultured for 24 hrs before media change and selection with hygromycin B (50 µg/ml) for a further 72 hrs. The surviving cells were trypsinised and subjected to limited dilution and incubated in 96-well plates for 10-14 days. Cells from wells containing only single colonies were expanded and genotyped to assess the mutational status of *DBF4* or *DRF1*. For screening, cells were washed with PBS and lysed in 50 μl of lysis buffer (10 mM Tris-HCl pH 7.5, 10 mM EDTA, 10 mM NaCl, 0.5% (w/v) N-Lauryl sarcosine, 10 μg/ml proteinase K and 20 μg/ml glycogen) for 2 hrs at 60°C. Genomic DNA was precipitated by adding 3x volumes of 150 mM NaCl in 96% (v/v) EtOH before mixing and incubation at room temperature for 30 min. DNA was pelleted by centrifugation (15 min, 800 x g), washed with 70% (v/v) EtOH and re-pelleted prior to air drying and resuspension in 100 μl TE buffer (10 mM Tris-HCl pH 7.5, 1 mM EDTA). 5 μl genomic DNA were used in PCR reactions using Taq DNA Polymerase and screening primers (DBF4-fwd: 5’-ACTTTGTTCTCTTCTAGCGAGTTG-3’, DBF4-rev: 5’-CGCCATCCCTAAATACAAGGGT-3’; DRF1-fwd: 5’-GGCTCTTAGGCTTTGGCAGA-3’, DRF1-rev: 5’-GTCAGCATACACCCAAGGGG-3’). PCR products were purified with MACHEREY-NAGEL NucleoSpin® Gel and PCR clean-up kit and sequenced using DBF4-fwd or DRF1-fwd primer by Eurofins genomics. Genome editing was accessed by manual analysis of the Sanger sequencing reads.

### Protein samples preparation

For whole cell extracts cells were harvested, washed with PBS, and resuspended in 1 volume of 20% (v/v) Trichloroacetic acid (TCA). Samples were mixed and 2 volumes of 5% TCA before centrifugation at 3000 rpm for 10 min. Pellets were resuspended in appropriate volumes of 2x Laemmli buffer, pH of the extracts was neutralised using 1 M Tris base, and samples were denatured for 5 min at 95°C. Proteins were separated by SDS-PAGE.

For chromatin fractionation MCF10A cells were seeded at 150,000 per well in 6-well plates and following treatment with indicated reagents, cells were harvested, washed once with PBS and lysed in CSK buffer (10 mM PIPES pH 6.8; 300 mM sucrose; 100 mM NaCl; 1.5 mM MgCl_2_; 0.5% (v/v) Triton X-100; 1 mM ATP; 1 mM DTT; 1 mM sodium orthovanadate; 2 mM N-ethylmaleimide; Phosphatase Inhibitor Cocktail I and Protease Inhibitor Cocktail III (ThermoFisher)) for 10 min on ice. Samples were centrifuged at 1000 x g for 5 min at 4°C, and the supernatant was transferred to a new reaction tube (soluble fraction). The pellet was washed using CSK buffer (2x volume of soluble fraction), centrifuged at 1000 x g for 5 min at 4°C and the supernatant was discarded. Pellets were resuspended in CSK buffer containing benzonase (125 U/ml) and incubated for 30 min on ice. Samples were denatured for 5 min at 95°C in 1x Laemmli buffer (Chromatin-enriched fraction). Protein concentration of the soluble fraction was determined by Bradford assay, and the required amount of protein was denatured for 5 min at 95°C in 1x Laemmli buffer.

### Immunoblotting

Proteins were transferred onto 0.2 µm pore size nitrocellulose membranes using a wet blot transfer system (Biorad). Proteins on membranes were stained with fast green (0.0001% (w/v) fast green in 0.1% (v/v) acetic acid) for 5 min as a total protein stain (TPS) and were analysed on the Odyssey infrared imaging system at 680 nm (LI-COR Biosciences). Membranes were de-stained with 0.1 M NaOH in 30% (v/v) methanol for 10 min and washed three times in ddH2O for 5 min at room temperature. Membranes were blocked in 3% (w/v) skim milk (Sigma) in TBS-T (20 mM Tris-HCl pH 7.5, 150 mM NaCl, 0.05% (v/v) Tween-20) for 1 hr at room temperature. Membranes were incubated in primary antibody diluted in blocking buffer overnight at 4°C followed by three washes in TBS-T for 10 min each. Secondary antibodies were diluted in blocking buffer and membranes were incubated for 1 hr at room temperature (protected from light) followed by three washes in TBS-T for 10 min at room temperature. Signals were acquired using the Odyssey infrared imaging system and analysed using Image Studio 2.0.38 and Empiria software 1.3.0.83 (LI-COR).

Primary antibodies were diluted in 3% skim milk/TBS-T: CHK1 (sc8408; Santa Cruz Biotech; 1:1000), CDC7 (DCS-342; MBL; 1:1000), and MCM2 (in-house: 1:3000) or 1% BSA/TBS-T: pT1989 ATR (GTX128145; GeneTex; 1:1000), pS345 CHK1 (2348, CST, 1:1000), pS139 H2AX (9718; CST, 1:1000), pS4/8 RPA32 (1:1000, A300-254A; Bethyl Laboratories) and pS40/S41 MCM2 (in house; 1:3000). IRDye secondary antibodies (LI-COR): 800CW goat anti-rabbit (926-32211; LI-COR Biosciences; 1:10,000) and 800CW goat anti-mouse (926-32210; LI-COR Biosciences; 1:10,000) were diluted in the same buffer as the primary antibody.

### DNA-mediated chromatin pull-down (Dm-ChP)

For analysis of proteins that were associated with nascent DNA, cells were plated at 9×10^6^ cells in 150 mm plates. Following treatment, cells were labelled with 10 µM EdU for 30 min, processed and then analysed using the DNA-mediated chromatin pull-down (Dm-ChP) technique^49^.

On the day before harvesting, streptavidin agarose beads were prepared (100 µl per sample, 50% slurry) by washing the required amount of beads three times with 1 ml of Wash buffer (10 mM Tris-HCl pH 8.0, 140 mM NaCl, 0.5 mM DTT) and centrifugation for 2 min, at 1,200 rpm and 4°C. Then, beads were blocked overnight at 4°C (under constant rotation) with 1 ml of blocking buffer (0.5 mg/ml BSA and 0.4 mg/ml pre-sheared salmon sperm DNA in RIPA buffer (10 mM Tris-HCl pH 8.0, 140 mM NaCl, 0.1% (w/v) sodium deoxycholate, 0.1% (v/v) SDS, 1% (v/v) Triton X-100 with Phosphatase Inhibitor Cocktail I and Protease Inhibitor Cocktail III)) to minimises non-specific binding. Beads were centrifuged for 2 min at 1200 rpm and 4°C, the supernatant was discarded and the beads were transferred to a new tube. The beads were washed twice with 1 ml Wash buffer and centrifuge for 2 min at 1,200 rpm and 4°C. For the last wash beads were resuspended in 500 μl of Wash buffer and transferred to a new tube and stored at 4°C until cell lysates were prepared. Before use beads were washed one last time (2 min at 1200 rpm and 4°C) and resuspended in an appropriate volume RIPA buffer to achieve 50% slurry.

Following the labelling with EdU, cells were washed twice with PBS before fixing the cells for 10 min at room temperature using serum-free media containing 1% PFA. To quench the PFA, 0.125 M Glycine was added to the plates and cells were incubated for 10 min at room temperature on a shaker. The media was discarded and plates were washed three times with 10 ml of ice-cold PBS. In the following cells were scraped of the plates using a cell scraper and transferred into a 15 ml falcon tube. The samples were centrifuged (1200 rpm, 5 min, 4°C), supernatant was removed and cells were permeabilised with the addition of 1 ml 0.1% (v/v) Triton-X100 in PBS with Phosphatase Inhibitor Cocktail I and Protease Inhibitor Cocktail III (ThermoFisher) for 10 min on ice. Triton/PBS solution was removed (1200 rpm, 5 min, 4°C), washed once with ice-cold PBS before performing a click reaction by gently resuspending the samples in 1 ml of click reaction mix (10 mM Sodium-L-ascorbate, 0.1 mM Biotin-TEG azide, 2 mM CuSO_4_, in PBS) and incubating them in the dark for 30 min at room temperature. After the incubation, 10 ml of PBS-T (0.5% (v/v) Tween-20; 1% BSA (w/v) in PBS) were added and samples were incubated for an additional 10 min to remove excess copper and azide. Samples were washed in PBS (1200 rpm, 5 min, 4°C) and cell pellets were resuspended in 1.2 ml of Cytoplasmic Lysis buffer (50 mM HEPES pH 7.8, 150 mM NaCl, 1.5 mM MgCl_2_, 0.5% (v/v) NP-40, 0.25% (v/v) Triton X-100, 10% (v/v) glycerol) and transferred to 1.5 ml Eppendorf tubes. Samples were rotated for 10 min at room temperature, before being centrifuged at 1300 rpm for 5 min. The supernatant was collected as the soluble fraction. The pellets were washed with 1.2 ml Wash buffer and rotated for 10 min at room temperature before being centrifuged at 1300 rpm for 5 min at room temperature. The pellets were resuspended in 1.2 ml RIPA buffer (10 mM Tris-HCl pH 8.0, 140 mM NaCl, 0.1% (w/v) sodium deoxycholate, 0.1% (v/v) SDS, 1% (v/v) Triton X-100 with Phosphatase Inhibitor Cocktail I and Protease Inhibitor Cocktail III (ThermoFisher)) and incubated under constant rotation for 5 min at room temperature. In the next step, samples were sonicated using a digital sonifier (Branson) according to the following conditions: 40% amplitude, 1 second pulse on, 10 seconds pulse off, 10 cycles. This sonication was repeated 6 times for each sample to fragment the labelled DNA. After sonication, samples were centrifuged at 12,000 x g for 10 min at 4°C and supernatant was transferred into a new tube. The protein concentration of the samples was quantified using a BCA assay kit (ThermoFisher). Equivalent amounts of protein were removed for each sample and RIPA buffer was added to a final volume of 1 ml. Additionally, 15-20 µg of protein were removed as input sample for later use. 100 µl of pre-blocked streptavidin agarose beads were added to each sample before incubation overnight at 4°C and constant rotation (12 rpm). After the incubation, the immunoprecipitation (IP) samples were centrifuged at 1200 rpm for 2 min and supernatant was removed as flowthrough. The samples (beads) were wash 6 times with 1 ml of wash buffer and were then resuspended in 70 μl of RIPA buffer with 1x Laemmli buffer. Samples were incubated at 95°C for 5 min and the IP eluate was carefully removed using a Hamilton syringe. Samples were analysed via SDS-PAGE and Western blot analysis.

### Replication timing analysis

20,000,000 exponentially growing cells were incubated, protected from light, with 50 μM BrdU (Abcam, #142567) at 37°C for 90 min. Cells were then fixed in 75% final concentration cold EtOH and stored at −20°C. BrdU labelled cells were incubated with 80 μg/mL Propidium Iodide (Invitrogen, P3566) and with 0.4 mg/ml RNaseA (Roche, 10109169001) for 1 hr at room temperature. 150,000 cells were sorted in early (S1) and late (S2) S-phase fractions using a Fluorescence Activated Cell Sorting system (FACS Aria Fusion, BD) in Lysis Buffer (50 mM Tris pH 8, 10 mM EDTA, 0.5% SDS, 300 mM NaCl). DNA from each fraction was extracted using Proteinase K treatment (200 µg/ml, Thermo Scientific, EO0491) followed by phenol-chloroform extraction and sonicated to a size of 500-1,000 bp, as previously described^43^.

Immunoprecipitation was performed using the IP star robot at 4°C (indirect 200 µl method, SX-8G IP-Star® Compact Automated System, Diagenode) with an anti-BrdU antibody (10 μg, purified mouse Anti-BrdU, BD Biosciences, #347580). Denatured DNA was incubated 5 hrs with anti-BrdU antibodies in IP buffer (10 mM Tris pH 8, 1 mM EDTA, 150 mM NaCl, 0.5% Triton X-100, 7 mM NaOH) followed by an incubation of 5 hrs with Dynabeads Protein G (Invitrogen, 10004D). Beads were then washed with Wash Buffer (20 mM Tris pH 8.0, 2 mM EDTA, 250 mM NaCl, 1% Triton X-100). Reversion was performed at 37°C for 2 hrs with a solution containing 1% SDS and 0.5 mg Proteinase K followed, after the bead’s removal, by an incubation at 65°C for 6 hrs in the same solution.

Immunoprecipitated BrdU-labelled DNA fragments were extracted with phenol-chloroform and precipitated with cold ethanol. Control quantitative PCRs (qPCRs) were performed using oligonucleotides specific of mitochondrial DNA, early (BMP1 gene) or late (DPPA2 gene) replicating regions^43^. Whole genome amplification was performed using SeqPlex^tm^ Enhanced DNA Amplification kit as described by the manufacturer (Sigma-Aldrich, SEQXE). Amplified DNA was purified using PCR purification product kit as described by the manufacturer (Macherey-Nagel, 740609.50). DNA amount was measured using a Nanodrop. Quantitative PCRs using the oligonucleotides described above were performed to check whether the ratio between early and late replication regions was still maintained after amplification. Early and late nascent DNA fractions were labelled with Cy3-ULS and Cy5-ULS, respectively, using the ULS arrayCGH labelling Kit (Kreatech, EA-005). Same amounts of early and late-labelled DNA were loaded on human DNA microarrays (SurePrint G3 Human CGH arrays, Agilent Technologies, G4449A). Hybridization was performed at 65°C as previously described^43^. The following day, microarrays were scanned using an Agilent C-scanner with Feature Extraction 9.1 software (Agilent technologies). The START-R suite was used to analyse the data^44^. Differential analysis of two experiments, each composed of two technical replicates, were performed with START-R analyzer and visualized with START-R viewer.

### Genomic studies of Advanced and Delayed replication timing domains

For each experiment, START-R Analyzer^38^ generated segmentation bed files corresponding to early, mid, late, TTR, advanced and delayed replicating domains. We also generated a random sample on the Galaxy website from the initial file containing the genomic coordinates of all advanced and delayed replicating domains detected after CDC7 inhibition and DBF4 loss, using successive rounds of Galaxy bedtools ShuffleBed randomly redistribute intervals in a genome (Galaxy Version 2.30.0) to obtained a random sample of 50,700 genomic regions.

The coverage of the different replicating domains and the random sample with large genes (>400kb), constitutive origins, CpG islands and putative G4 were done with the Coverage of a set of intervals on second set of intervals software (Galaxy Version 1.0.0). Boxplots illustrating differences in these coverages were generated. The constitutive origins and the putative G4 files (hg19 genome assembly) were taken from Picard et al., 2014^39^ and converted with LiftOver to hg18 genome assembly. Position of genes came from the UCSC table browser RefSeq Genes database without duplicates and with hg18 genome assembly. Large genes regions (>400kb) were extracted from the aforementioned gene database. The CpG islands file came from UCSC browser (hg19 assembly) and was converted with LiftOver to hg18 genome assembly. Data are deposited in GEO with accession number GSE248981.

### Fluorescence microscopy

MCF10A cells were seeded at a density of 130,000 cells per well in a 6-well plate on poly-L-lysine coated coverslip. Following drug treatment, coverslips were washed once with PBS and cells were fixed in PBS containing 4% (w/v) PFA for 10 min at room temperature. Following the fixation step, samples were washed three times with PBS to remove residual PFA. Cells were permeabilized with PBS-TX (0.1% (v/v) Triton X-100 in PBS) for 20 min at room temperature followed by incubation for 30 min in blocking buffer (10% (v/v) FBS, 0.5% (w/v) BSA in PBS-TX) at room temperature. Cells were incubated for 1 hr at 37°C with mouse anti-beta-Tubulin (#05-661; Millipore; 1:1500) primary antibody diluted in blocking buffer. Following three washes with PBS-TX, coverslips were incubated for 1 hr at 37°C with goat anti-mouse AlexaFluor 488 (A11001; ThermoFisher; 1:500) secondary antibody diluted in blocking buffer, which additionally contained DAPI (1:600 dilution in PBS of 0.5 mg/ml stock) to stain nuclei. Cover slips were washed three times in PBS-TX, once in PBS and dipped in ddH2O before being mounted onto slides using SlowFade Gold Antifade Reagent (ThermoFisher).

Microscopy for micronuclei detection was performed using IX71 Olympus microscope using a 40X objective lens or 60X oil-immersion objective as indicated. The number of micronuclei per cell were manually counted using DAPI staining. Beta-Tubulin staining was used as cytoplasmic staining to assist in assigning micronuclei to specific cells. A minimum of 275 cells per sample of four independent experiments were analysed. After quantification and analysis, the brightness of representative images was adjusted to the same extent for all samples in an experiment to aid visualisation in figures.

### DNA combing

Cells were seeded at a density of 300,000 cells per well in a 6-well plate and allowed to recover overnight. To assess the replication fork speed in MCF10A cells in the presence of CDC7i or upon mutation of DBF4, cells were treated with DMSO or 10 µM XL413 for 24 hrs prior to labelling with 25 µM IdU for 30 min. Following the incubation, the media was exchanged and cells were washed three times with pre-warmed culture media followed by treatment with DMSO or 10 µM XL413 and labelling with 250 µM CldU for 30 min. Immediately following the CldU pulse 1 mM thymidine was added to block further CldU incorporation. Cells were harvested and washed with ice-cold PBS (5 min, 2000 rpm, 4°C). The cell pellets were resuspended in 110 µl ice-cold PBS, counted using Trypan blue exclusion, and cell number was adjusted to 1.5×10^6^ cells/ml. A 1.5% (w/v) low melting point agarose solution (in PBS) was prepared and stored at 45°C until required. 100 µl of the prepared cell suspension (=1.5 x10^5^ cells) were incubated at 45°C for 1 min before gently mixing it with 100 µl of the agarose solution and immediately transferring the mixture into two plug moulds. The agarose plugs were allowed to set for 5 min at room temperature and then transferred to 4°C for 30 min to solidify. For cell lysis, plugs were incubated in 0.3 ml proteinase K buffer (10 mM Tris-HCl pH 7.5, 50 mM EDTA, 1% (w/v) Sarkosyl, 2 mg/ml Proteinase K) overnight at 50°C. The next day, plugs were transferred into 10 ml TE50 buffer (10 mM Tris-HCl pH 7.5, 50 mM EDTA) and stored at 4°C until further use. To melt the agarose plugs, each plug was washed two times in 10 ml TE-Wash buffer (10 mM Tris-HCl pH 7.5, 1 mM EDTA) for 1 hr with rotation at room temperature, before being transferred into 2 ml MES buffer (0.5 M MES hydrate pH 5.5) for 1 hr at 68°C. The resulting solution was cooled down to 42°C before adding 2 U of β-agarase (NEB), gently mixing the solution by end-over-tube inversion, and incubating it overnight at 42°C. Before combing, the samples were incubated at 68°C for 10 min, cooled down to room temperature and transferred into Teflon reservoirs. Silanized coverslips were inserted into the combing apparatus, incubated in the DNA solution for 10 min before being automatically withdrawn by the DNA combing apparatus at a speed of 300 μm/s. Following the combing, slides were mounted onto glass slides and DNA was cross-linked to the slides by incubating them for 4 hrs at 60°C followed by storage at −20°C overnight. Slides were allowed to reach room temperature and combed DNA was denatured for 15 min in 0.05 M NaOH. Slides were washed three times in ice-cold PBS for 1 min to neutralise the NaOH and then dehydrated by incubating the slides in 70%, 90% and 100% EtOH in succession for 3 min each. After allowing the slides to dry at room temperature (in the dark), samples were blocked for 15 min in blocking buffer (1% (w/v) BSA/PBS). Next, slides were taken through a series of 30 min primary and then secondary antibody incubations in blocking buffer at room temperature, with three PBS washes in between each antibody incubation. The antibodies were used in the following order and concentrations: BrdU (BU1/75) rat monoclonal antibody (MA1-82088; ThermoFisher; 1:100) and Chicken anti-rat Alexa Fluor 488 (A21470, ThermoFisher; 1:300) to detect incorporated CldU. Anti-BrdU (B44) IgG1 mouse monoclonal (347580; BD biosciences; 1:100) and Goat anti-mouse IgG1 Alexa Fluor 546 (A21123, ThermoFisher; 1:300) to detect incorporated IdU, anti-ssDNA poly dT (mab3034, Millipore, 1:100) and Goat anti-mouse IgG2a Alexa Fluor 647 (A21241, ThermoFisher; 1:300). After the final antibody, coverslips were washed two times in PBS, once in ddH2O and allowed to dry completely (protected from light). Coverslips were mounted onto the silanized coverslips using SlowFade Gold Antifade Reagent (ThermoFisher). Images were captured with an IX71-Olympus microscope and 60X oil-immersion objective and analysis was performed manually in ImageJ Fiji software.

### Flow cytometry

Cell cycle distribution and rate of DNA synthesis were analysed by plating MCF10A cells at 150,000 per well in 6-well plates. Following treatment with indicated inhibitors or DMSO, nascent DNA was labelled by incubating cells with 10 μM EdU for 30 min at 37°C prior to harvesting. Cells were washed with PBS, which if not specified otherwise, involves centrifugation at 400 x g for 5 min at 4°C and the removal of the supernatant from the cell pellet. Samples were fixed by resuspension in 0.3 ml PBS and dropwise addition of 0.7 ml 100% ethanol while vortexing followed by incubation for at least 1 hr at −20°C. After a PBS wash, cells were washed with 1% (w/v) BSA/PBS and incorporated EdU was labelled by incubating cells in click reaction buffer (PBS containing 10 mM Sodium-L-ascorbate, 2 mM Copper-II-Sulfate and 10 µM 6-Carboxyfluorescine-TEG-azide) for 30 min at room temperature protected from light. Cells were then incubated in PBS containing 1% (w/v) BSA and 0.5% (v/v) Tween-20 for 5 min at room temperature followed by an additional wash with PBS. DNA was stained with 1 μg/ml DAPI in 1% (w/v) BSA/PBS and to reduce RNA interference 0.5 μg/ml RNase A was added to each sample until measurement. Fluorescence intensity data for DAPI (405_450_50 nm) and 6-Carboxyfluorescine (488_530_30 nm) were acquired for 10,000 single cells on a BD FACS Canto II and analysed using FlowJo 10.0.7 software. To perform analysis of fluorescence intensity proportional to EdU incorporation for individual cells, gates were applied on DAPI-EdU biparametric dot plots to select for EdU-positive (late) S-phase cells using FlowJo. Fluorescence intensity (488_530_30-H) values per cell were exported and plotted as scatter dot plots using GraphPad Prism 10.0.2.

Detection of pS139 H2AX via flow cytometry was performed by plating 400.000 MCF10A cells on 6 cm plates and treating them as described for the respective experiments. Following the treatment, MCF10A cells were detached using trypsin/EDTA, and washed once in ice-cold PBS (450 g, 5 min, 4°C). The washing steps were performed with the same centrifuge settings if not described otherwise. Next, cells were incubated in 300 µl 0.2% (v/v) Triton X-100/PBS for 10 min on ice followed by a washing step with PBS to remove excess Triton X-100 (450 g, 5 min, 4°C). Cell pellets were resuspended in 0.5 ml 1% (w/v) PFA/PBS and incubated for 10 min at room temperature. Cells were first washed in blocking buffer (1% (w/v) BSA/PBS) before being incubated for 30 min at 4°C in blocking buffer. The blocking buffer was removed (450 g, 5 min, 4°C), and cells were permeabilised by incubation in 0.05% (w/v) saponin buffer for 10 min on ice. In the following, samples were sequentially incubated with a primary antibody against pSer139 histone H2AX (1:500, #9718; CST) for 2 hrs, followed by washing in saponin buffer, and then incubation with a secondary antibody (1:250, Donkey anti-rabbit Alexa Fluor 488; A21206; ThermoFisher) for 30 min in the dark. Following two washing steps, cells were resuspended in 0.5 ml blocking buffer containing 0.1 mg/mL RNase and 1 µg/mL DAPI and incubated for 30 min in the dark at room temperature before measurement at the flow cytometer. Measurements and analysis were performed with the tools and settings described for EdU/DAPI staining.

### siRNA transfections

MCF10A cells were seeded at 60,000 cells per well in a 6-well plate and transfected 24 hrs after plating. siRNA transfections were performed using JetPRIME transfection reagent (Polyplus Transfection). For each individual well, a pool of 4 siRNAs (50 or 100 nM final concentration, as indicated) were prepared by mixing with JetPrime buffer (200 µl) and JetPrime reagent (4 µl) and the mixture was incubated for 15 min at room temperature prior to dropwise addition to the cells. Following a 5 hrs incubation at 37°C, 5% CO_2_ the culture media was exchanged for fresh, incubator-equilibrated media and cells were returned to the incubator. In this study, treatment of siRNA transfected cells was performed 48 hrs after transfection. Protein depletion was confirmed by qPCR. The following siRNA sequences were used: siCtrl: 5’-AGUACUGCUUACGAUACGG-3’; siDBF4_1: 5’-GAACACACAUUAAGUGAAA-3’; siDBF4_2: 5’-GAGCAGAAUUUCCUGUAUA-3’; siDBF4_3: 5’-GCACAAACCUUGGGUCGAA-3’; siDBF4_4: 5’-CCAAACAGAUGGCGAUAAG-3’; siDRF1_1: 5’-GGAAACAUCGGCCAUGGUU-3’; siDRF1_2: 5’-GGAAACCCGUUGACUCGGU-3’; siDRF1_3: 5’-AAACAUCGGCCAUGGUUGA-3’; siDRF1_4: 5’-GAGCGAACCGGGAAAGGGA-3’.

### Real-time qPCR

MCF10A EditR cells were transfected with siRNA as described in *siRNA transfections* 72 hrs before total RNA was extracted using the NucleoSpin RNA columns (Machery Nagel) according to manufacturer instructions. The concentration of the total RNA was determined using a NanoDrop. To generate cDNA 1 µg of RNA was used with random hexamer primers in the SuperScript First Strand Synthesis System (ThermoFisher). The resulting cDNA was diluted 1:10 and used for RT-qPCR using the following TaqMan expression assays: Hs00272696_m1 (DBF4), Hs010691951_m1 (DRF1), Hs99999901_s1 (18S). The relative mRNA levels were normalised to the 18S rRNA expression and calculated using the comparative Ct method.

## SUPPLEMENTARY INFORMATION

**Figure S1, related to Fig 1:**
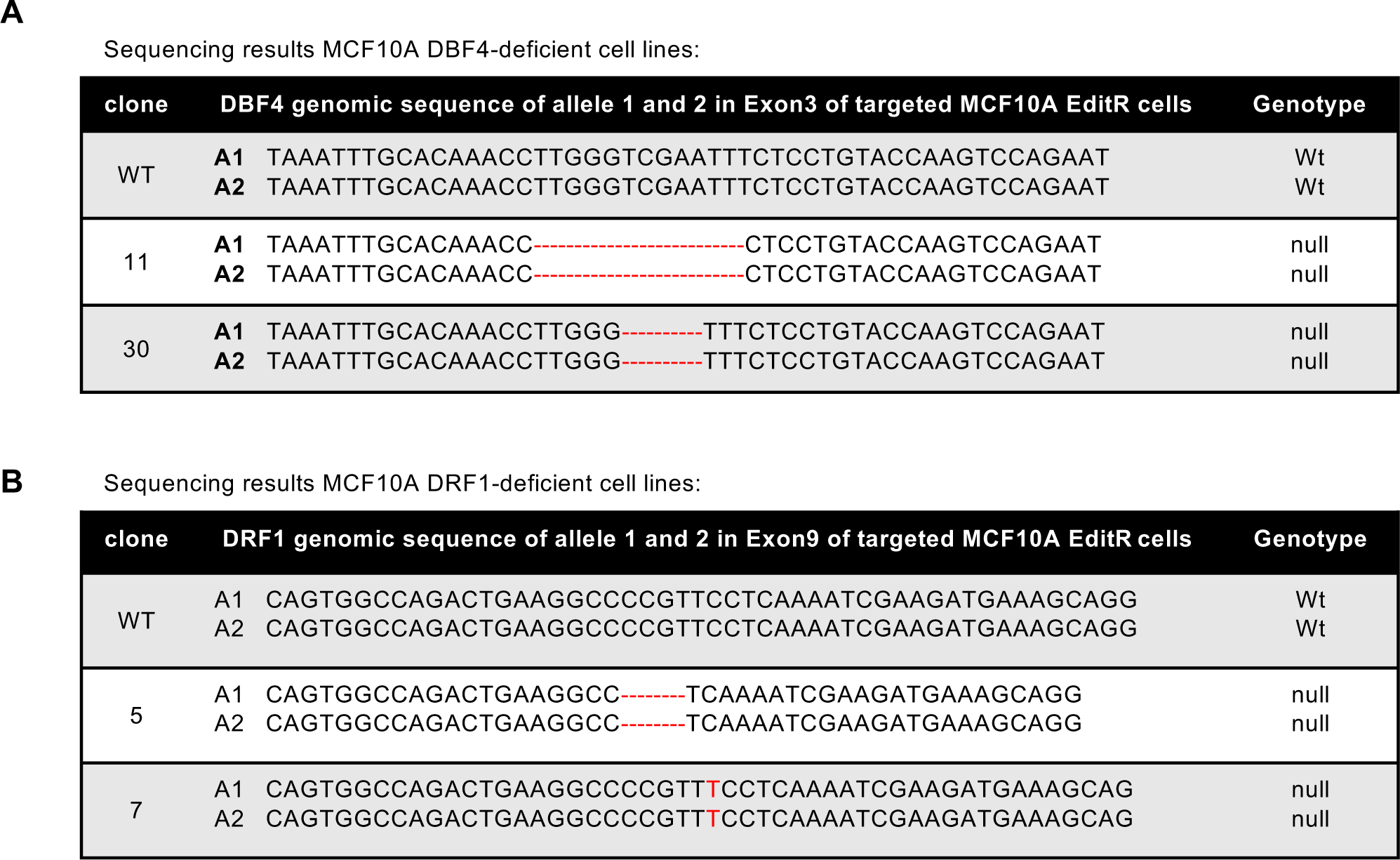
Characterisation of MCF10A DBF4- and DRF1-deficient cells. **A-B.** Sequencing results for MCF10A DBF4-11 and −30 (**A**) and MCF10A DRF1-5 and −7 (**B**). Original sequence (WT) around Cas9 cut site is displayed alongside sequences of individual clones.

**Figure S2, related to Fig 2:**
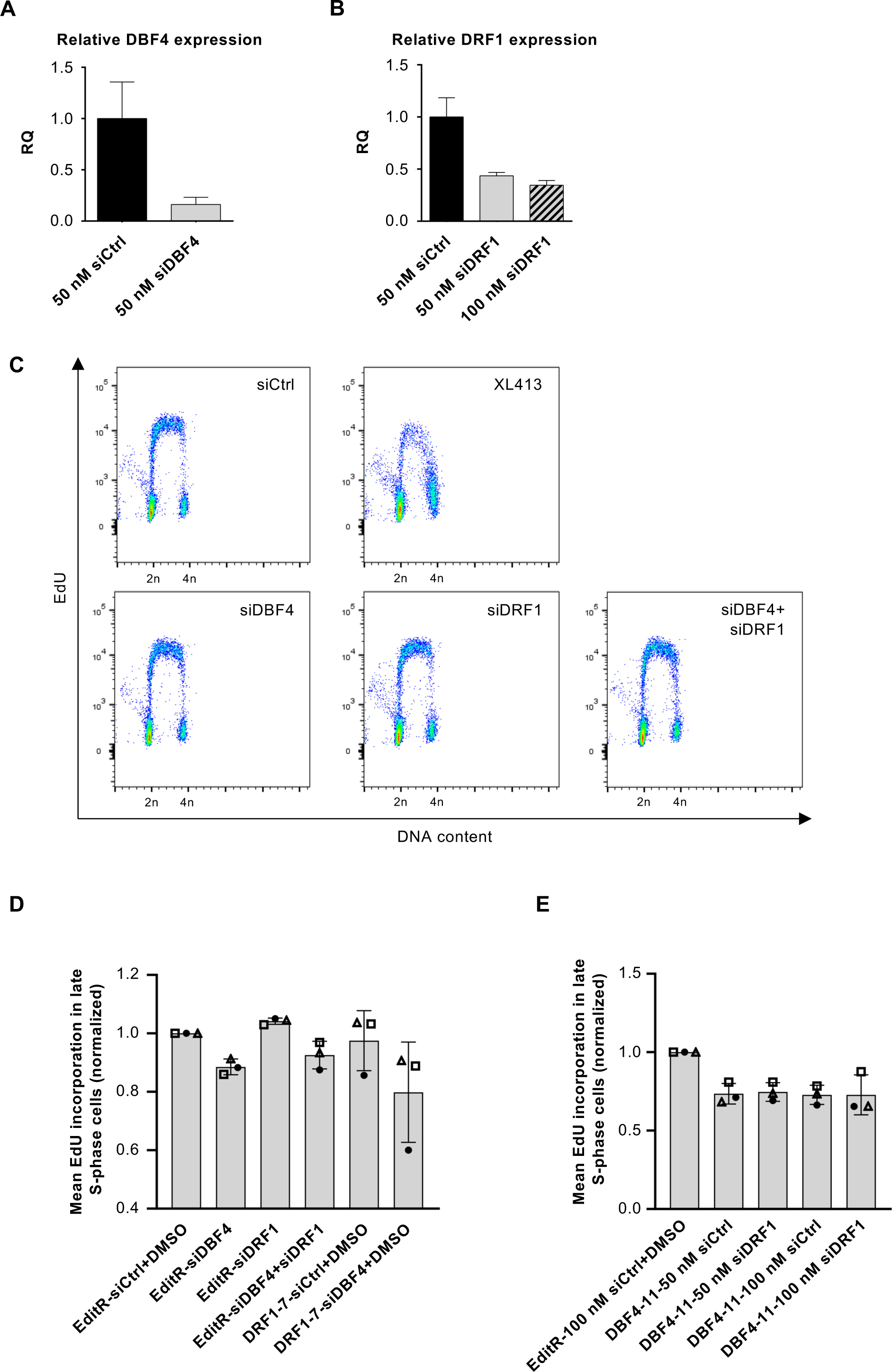
DBF4 depletion partially reduces DNA replication rate in MCF10A cells. **A-B.** MCF10A EditR cells were transfected with siCtrl, siDBF4 (**A**) or siDRF1 (**B**) at indicated concentrations over 72 hrs. DBF4 and DRF1 mRNA levels were assessed using Realtime-qPCR. Data are representative of one independent experiment performed in technical triplicates. **C.** MCF10A EditR were transfected with 50 nM of indicated siRNAs for 48 hrs followed by treatment with 10 µM XL413 or DMSO for 24 hrs. For flow cytometry analysis, cells were labelled with 10 µM EdU 30 min prior to harvest. Representative images from one of three independent experiments are shown. **D.** Analysis of fluorescence intensity, proportional to EdU incorporation, in late S-phase cells described in **C** and including additional samples of DRF1-7 cells transfected with 50 nM siCtrl or siDBF4 of the same experiment. Mean fluorescence intensity in late S-phase cells of three independent experiments was expressed as a ratio relative to MCF10A EditR cells transfected with siCtrl. Experiments are represented with different symbols, and columns are displayed as mean ± SDs. **E.** Analysis of fluorescence intensity, proportional to EdU incorporation, in MCF10A EditR and DBF4-11 cells transfected and treated as indicated in late S-phase of three independent experiments. Mean fluorescence intensity in was expressed as a ratio relative to MCF10A EditR cells transfected with siCtrl. Experiments are represented with different symbols, and columns are displayed as mean ± SDs.

**Figure S3, related to Fig 3:**
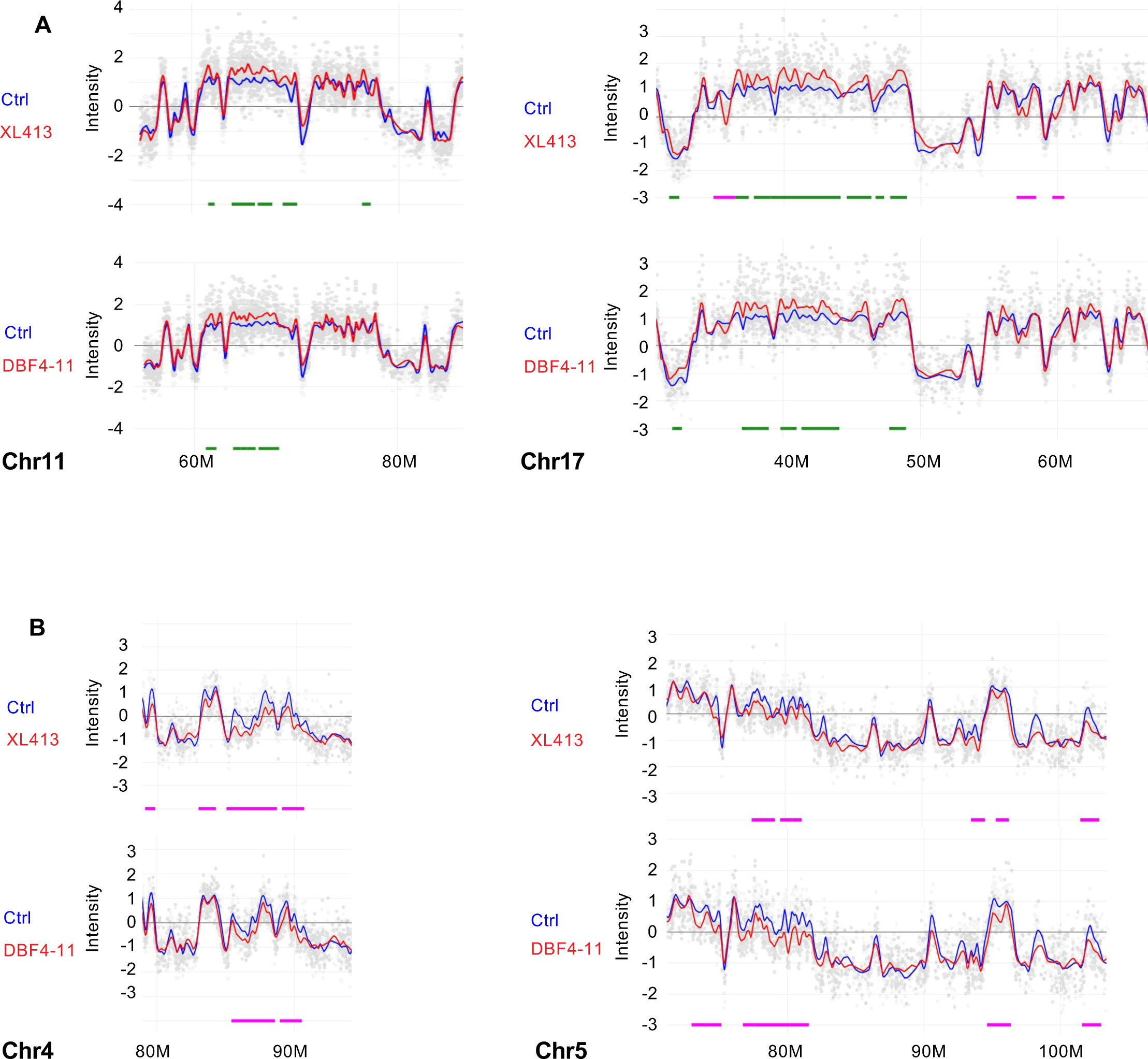
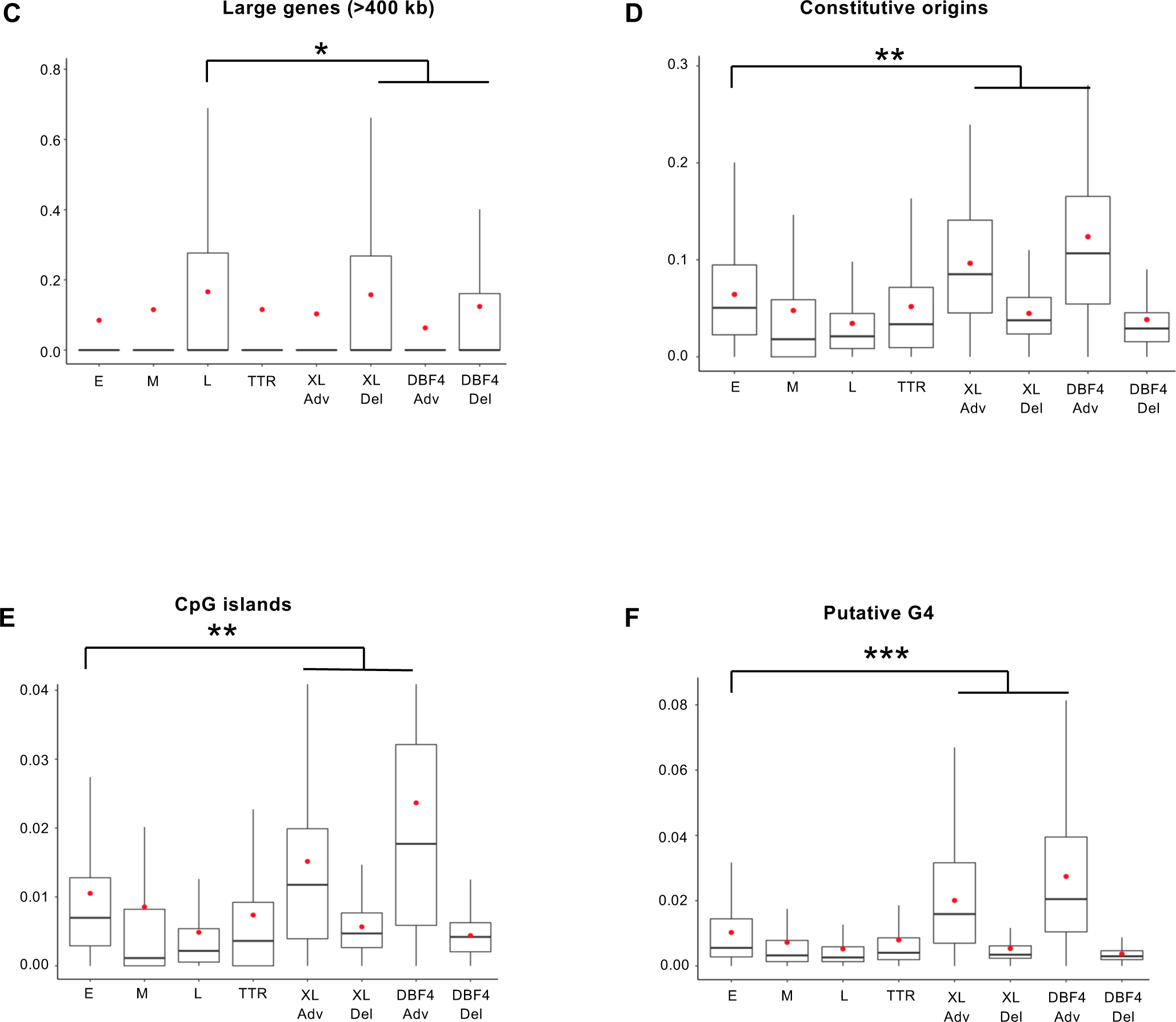
Regions with changed replication timing in MCF10A cells treated with XL413 or in DBF4-deficient cells share similar properties. **A.** Part of chromosomes 11 and 17 replication timing profiles with mostly advanced regions upon CDC7 inhibition with XL413 (top profiles) or in DBF4-deficient cells (bottom profiles). Blue line represents MCF10A EditR cells treated with DMSO, the red one cells treated with 10 µM XL413 or DBF4-deficient cells (DBF4-11). Chromosome coordinates are indicated below the profile in megabases. Differences in replication timing are marked below the profiles with advanced regions in green and delayed regions in magenta. Data is representative of two replicates of four independent experiments. **B.** Part of chromosomes 4 and 5 replication timing profiles with delayed regions upon CDC7 inhibition with XL413 (top profiles) or in DBF4-deficient cells (bottom profiles). Analysis was performed and graphs were generated as described in **A**. **C-F.** Coverage of large genes (>400 kb; **C**), constitutive origins (**D**), CpG islands (**E**), and regions rich in putative G4 sequences (**F**) in replication timing changing regions of MCF10A EditR and DBF4-deficient cells treated with 10 µM XL413 or DMSO for 24 hrs. Results were compared to the coverage of these factors in Early, Mid, and Late replicating regions or Timing Transition Regions (TTR) of MCF10A EditR cells. Advanced regions (Adv) and Delayed regions (Del) are displayed separately. The box plots show the dispersion of the data with a range from the 25^th^ to 75^th^ percentile, the sample median represented by the line inside the box and the mean by a red dot. Significance of the differences was estimated with a Wilcoxon test (* p<10^-3^, ** p<7.5 10^-5^ and *** p<7.3 10^-13^).

**Figure S4, related to Fig 5:**
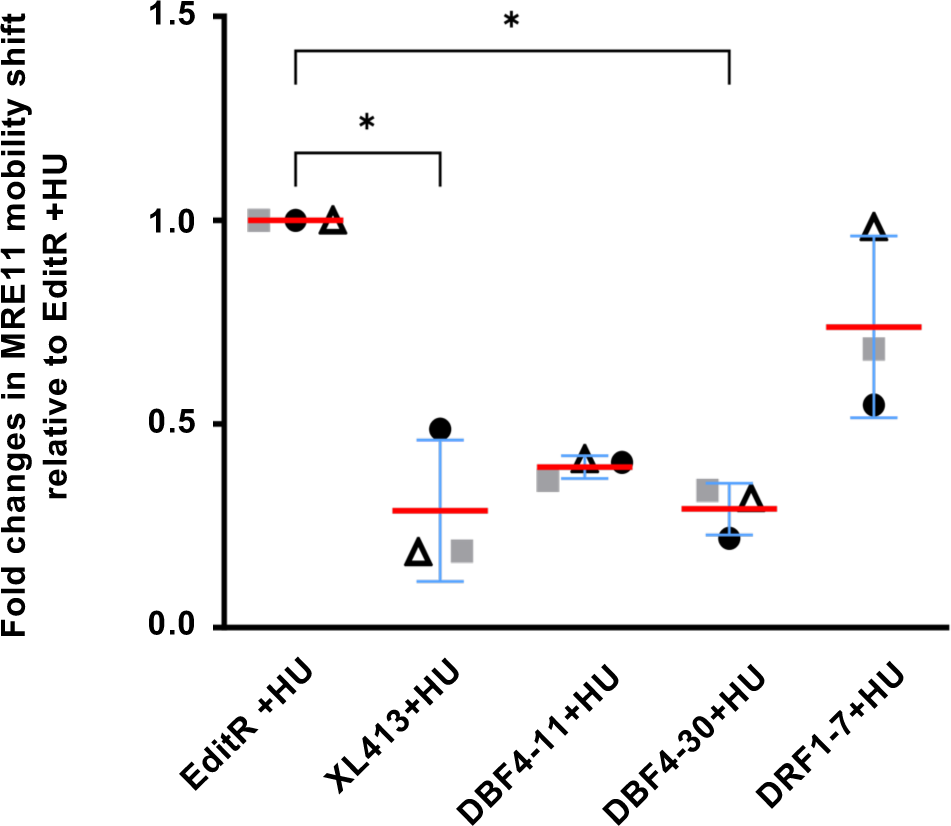
MRE11 mobility shift is primarily mediated by DBF4. Quantification of MRE11 mobility shift during electrophoretic analyses as presented in Figure 5A. Data shows fold change in MRE11 mobility shift of HU-treated DBF4-11, DBF4-30, DRF1-7, and XL413-treated EditR samples relative to MCF10A EditR co-treated with DMSO and HU. Data was normalised to TPS to account for differences in loading. Data are presented as mean fold change ± SD. Statistical analysis was performed using Kruskal-Wallis ANOVA, with Dunn’s multiple comparison test (*<0.05).

